# LIVIA: a browser-based tool for assessing and visualizing predicted protein interactions

**DOI:** 10.64898/2026.05.01.721633

**Authors:** Ah-Ram Kim, Norbert Perrimon

## Abstract

As protein structure prediction tools become widely adopted across biology, there is a growing need for accessible methods to assess and visualize predicted protein-protein interactions (PPIs). Here we present LIVIA (Local Interaction Visualization and Analysis), a browser-based tool that computes local PPI confidence metrics across multiple prediction platforms, identifies predicted interface residues, embeds an interactive Mol* 3D viewer, and generates visualization scripts for ChimeraX and PyMOL. The tool automatically detects prediction formats; all parsing and computation occur locally on the user’s machine. LIVIA is freely available at https://flyark.github.io/LIVIA.

## 1 Introduction

Deep learning methods for predicting protein complex structures are now routinely used to study protein-protein interactions (PPIs) [*1–7*]. Beyond the predicted three-dimensional structure, these methods report confidence estimates for each residue and residue pair. For example, the Predicted Aligned Error (PAE) quantifies the confidence in the predicted relative position of any two residues, a quantity especially useful for evaluating predicted interfaces.

Given a prediction, two questions arise: whether the proteins interact and, if so, which residues form the interface. Recent local confidence metrics that focus on interface residues have proven more informative than global scores for both tasks, enabling more sensitive detection of small or flexible interfaces [*8–11*]. However, computing these metrics and visualizing the results typically requires platform-specific parsing code and custom scripts for tools such as ChimeraX [*12*] and PyMOL [*13*], an effort that scales poorly as the number of predictions grows.

Here we present LIVIA, a browser-based tool that bridges this gap by integrating local confidence scoring, interaction visualization, and script generation across multiple prediction platforms within a single interface. In addition, LIVIA retrieves and analyzes dimer predictions directly from the AlphaFold Database.

## 2 Description

### 2.1 Multi-platform input

LIVIA accepts prediction outputs from AlphaFold-Multimer, AlphaFold3, ColabFold, Boltz-1/2, Chai-1, and OpenFold3 (**Figure 1A**). Users upload prediction files in their native format, including ZIP archives. For example, a single ZIP file downloaded from the AlphaFold3 Server (which may contain multiple prediction jobs) can be loaded directly without extraction, and individual predictions can be selected and analyzed one at a time. The platform is automatically detected from filename patterns and data structures. LIVIA includes p53–MDM2 example predictions from each supported platform, loadable without file upload, to demonstrate consistent handling across formats.

**Figure 1.**
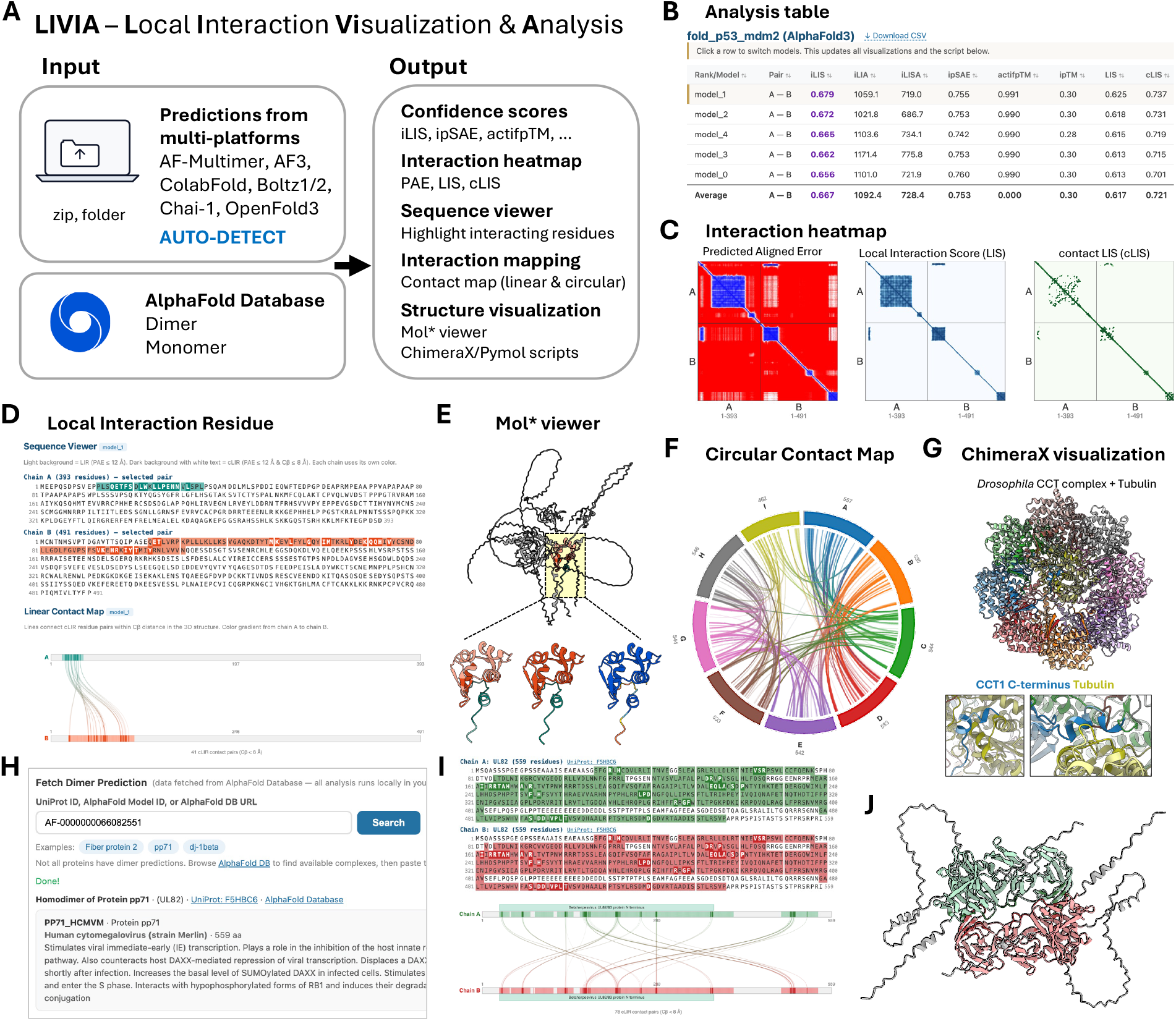
LIVIA overview and example outputs. **(A)** Input/output summary. LIVIA accepts prediction files (ZIP archives or folders) from AlphaFold-Multimer, AlphaFold3, ColabFold, Boltz-1/2, Chai-1, and OpenFold3 with automatic platform detection, and also retrieves dimer/monomer predictions directly from the AlphaFold Database. **(B)** Analysis table of per-model interface confidence metrics (iLIS, ipSAE, actifpTM, ipTM, LIS, and cLIS) shown for an AlphaFold3 p53–MDM2 prediction. **(C)** PAE, LIS, and cLIS heatmaps for the selected model. **(D)** Local Interaction Residue (LIR) panel: sequence viewer with LIR (light) and cLIR (dark) highlighting, and a linear contact map connecting inter-chain cLIR pairs. **(E)** Embedded Mol* 3D viewer of the selected model. The full structure is shown with non-interacting regions in gray and the LIR-containing domain highlighted by a yellow box. Three close-up renderings of the highlighted domain illustrate alternative coloring schemes: LIR and cLIR distinguished by shade, a single color per subunit, and pLDDT-based coloring restricted to confident regions. **(F, G)** Application to a large complex: AlphaFold3 prediction of the *Drosophila* TRiC/CCT chaperonin with α-tubulin (αTub67C; 9 chains, 4,795 residues; chains A–H are the eight CCT subunits, chain I is αTub67C). **(F)** Circular contact map revealing inter-chain cLIR contacts among all chains. **(G)** Structure rendered by the downloadable ChimeraX script (detailed view of the CCT1–αTub67C interface in **Figure S1D, E**). **(H–J)** AlphaFold Database Dimer analysis, illustrated with the human cytomegalovirus tegument protein pp71 (UL82; AFDB entry AF-0000000066082551). **(H)** Dimer search interface accepting UniProt ID, AlphaFold model ID, or AFDB URL, with a brief protein summary. **(I)** Sequence viewer and linear contact map for the fetched pp71 homodimer, with domain annotations retrieved from UniProt. **(J)** ChimeraX-rendered view of the fetched dimer. LIR are colored in the light chain color, cLIR in the dark chain color, and non-interacting residues are shown in gray.

### 2.2 Local interaction metrics and residues

For each chain pair, LIVIA computes interface confidence metrics from the AFM-LIS framework [*8, 11*] (**Figure 1B**). Briefly, the Local Interaction Score (LIS) averages PAE-derived local interaction probabilities across inter-chain residue pairs with PAE ≤ 12 Å; contact LIS (cLIS) restricts these pairs to those additionally in contact (Cβ ≤ 8 Å); and integrated LIS (iLIS) combines both into a single score. Interface residues are classified as Local Interaction Residues (LIR; PAE ≤ 12 Å) or contact-filtered LIR (cLIR; PAE ≤ 12 Å and Cβ ≤ 8 Å). For comparison with other commonly used scores, LIVIA computes ipSAE [*9*] and actifpTM [*10*] from the PAE matrix and inter-chain contacts, alongside the platform-reported ipTM for reference. For multi-subunit complexes, all pairwise metrics are computed and summarized in a score matrix overview.

### 2.3 Visualization and model comparison

LIVIA provides coordinated visualizations that update in response to model and chain-pair selection: PAE, LIS, and cLIS heatmaps (**Figure 1C**); a Local Interaction Residue panel combining a sequence viewer and a linear contact map with domain annotations (**Figure 1D**); residue-level intra-chain LIS and predicted local distance difference test (pLDDT) charts; and an embedded Mol* 3D viewer [*14*] (**Figure 1E**). The sequence viewer highlights LIR in light color and cLIR in dark color, and the 3D viewer displays only the LIR region by default with cLIR highlighted. Because prediction methods typically produce multiple models, LIVIA lists all models in a single table and overlays the selected model’s residue-level intra-chain LIS and pLDDT against the average ± standard deviation across models.

To support downstream analysis and reproducibility, LIVIA generates downloadable ChimeraX and PyMOL scripts that reflect the selected color scheme, and exports all computed metrics and LIR/cLIR assignments as CSV. In-tool documentation (per-page help sections and an About page) summarizes metric definitions and the recommended iLIS threshold.

### 2.4. Application to a multi-chain complex

To illustrate LIVIA on a large multi-chain complex, we examined the *Drosophila* TRiC/CCT (TCP-1 Ring Complex / Chaperonin Containing TCP-1) chaperonin. CCT is essential for cell proliferation and organ growth [*15, 16*] and is the archetypal chaperonin for folding cytoskeletal substrates including tubulin and actin [*17*]. CCT is an octameric ring of eight paralogous subunits (CCT1– CCT8) that in vivo stacks back-to-back into a hexadecameric folding chamber.

We predicted the *Drosophila* CCT complex with AlphaFold3 in apo form (8 subunits, 4,333 residues) and with its known substrate tubulin in both α (αTub67C; UniProt P06606) and β (βTub56D; UniProt A1ZBL0) isoforms. All three predictions were loaded into LIVIA directly as AlphaFold3 Server ZIP archives (167 / 208 / 215 MB). The substrate-bound predictions are markedly more confident than the apo prediction (**Figure S1A–C**). All three predictions recover the canonical TRiC ring topology (CCT1–CCT4–CCT2–CCT5–CCT7–CCT8–CCT6–CCT3), matching the consensus subunit arrangement [*17*].

For the αTub67C-bound prediction, LIVIA’s circular contact map of inter-chain cLIR contacts reveals connections across all 9 chains (**Figure 1F**). The ChimeraX rendering produced by the downloadable script shows αTub67C inside the cavity with CCT C-terminal tails contacting it (**Figure 1G**). Cryo-EM of human CCT/TRiC visualized the CCT2 and CCT6 C-tails directly engaging β-tubulin and proposed a CCT1–CCT2 C-terminal fulcrum for substrate repositioning [*17*]. In our AlphaFold3 predictions of *Drosophila* CCT with both αTub67C and βTub56D, the CCT1 C-terminal tail (residues 543–557) reproducibly contacts substrate, supporting this fulcrum hypothesis with a contact that averaging-based cryo-EM may miss due to the C-tail’s intrinsically disordered character (**Figure S1D–G**).

### 2.5 AlphaFold Database Dimer analysis

The AlphaFold Protein Structure Database (AFDB) [*18, 19*] recently expanded to include ∼1.7 million high-confidence homodimer structures [*20*], but AFDB provides only confidence scores without annotation of which residues form the predicted interaction interface. LIVIA’s AFDB Dimer analysis addresses this gap: users search by UniProt ID (if a dimer prediction is available), AlphaFold model ID, or AFDB URL, and LIVIA fetches the dimer prediction, computes all metrics including LIR/cLIR, and renders the full visualization pipeline (**Figure 1H–J**). It additionally retrieves the matching monomer predictions and overlays per-residue intra-chain LIS and pLDDT so users can identify residues that gain or lose folding confidence upon dimerization. The pipeline applies equally to heterodimers when they become available in future AFDB releases.

### 2.6 Additional modules

Beyond general interface analysis, LIVIA includes three additional modules that provide entry points into pre-computed interactomes and intra-chain interaction analysis. The FlyPredictome module searches *Drosophila* PPI predictions by gene name or gene pair [*11*]. The Ortholog Interactome module extends this lookup to ortholog-based predictions in human and other species within the FlyPredictome database. The Monomer Subdomain module extends LIVIA to intra-chain interactions such as autoinhibitory contacts by analyzing monomer structures from AFDB: it identifies subdomains and computes pairwise LIR/cLIR between them. Region detection parameters are user-adjustable, since automated subdomain identification can over-fragment in flexible proteins.

## 3 Conclusion

LIVIA combines local confidence scoring with interactive visualization across multiple prediction platforms, making predicted interface residues immediately accessible for individual predictions and large multi-chain complexes alike. By exposing residue-level interface annotations in a single interface, LIVIA lowers the barrier to mechanistic interrogation of individual predictions and to screening candidate interactors. It helps researchers who routinely generate or evaluate complex predictions and require rapid, reproducible interface assessment without writing custom scripts. LIVIA assumes the structures provided are predicted complexes; it does not perform docking or structure prediction itself. LIVIA’s modular input layer can be extended to accommodate new formats as new prediction methods emerge, and its AFDB Dimer module will apply directly to heterodimer entries when they become available.

## Availability mand Implementation

LIVIA is freely available at https://flyark.github.io/LIVIA with source code at https://github.com/flyark/LIVIA under the MIT license. The tool is implemented as a static web application: the browser downloads the page once, then performs all parsing, metric computation, and rendering using local CPU and memory, with no data uploaded to a server. LIVIA has been tested in Chrome, Firefox, and Safari. On an Apple M3 MAX MacBook, a 2-chain prediction (∼400 residues, 5 models; p53–MDM2) is analyzed in 1–4 s, while a 9-chain complex of 4,795 residues (*Drosophila* CCT + αTub67C, 5 models; AF3) completes in ∼30 s; performance is expected to be comparable on other current laptops with sufficient memory (peak ∼3.3 GB for the CCT case). Detailed metric definitions, the recommended iLIS threshold, and parameter descriptions are documented in the LIVIA About page and GitHub repository. A command-line Python tool (lis.py) for batch metric computation is available at https://github.com/flyark/AFM-LIS.

## Acknowledgments

This research was supported by NIH NIGMS P41 GM132087 and NIH NIAMS R01 AR057352. A.-R.K. was supported by the Postdoctoral Fellowship Program (Nurturing Next-generation Researchers) through the National Research Foundation of Korea (NRF) funded by the Ministry of Education (2021R1A6A3A14039622). N.P. is an investigator of the Howard Hughes Medical Institute.

During the development of LIVIA and the preparation of this manuscript, the authors used Anthropic’s Claude (Opus 4.6 and 4.7) to assist in generating code for the web application and to improve the readability and language of the text. After using this tool, the authors reviewed and edited the content as needed and take full responsibility for the publication.

This article is subject to HHMI’s Open Access to Publications policy. HHMI lab heads have previously granted a nonexclusive CC BY 4.0 license to the public and a sublicensable license to HHMI in their research articles. Pursuant to those licenses, the author-accepted manuscript of this article can be made freely available under a CC BY 4.0 license immediately upon publication.

## Supplementary Figure

**Figure S1.**
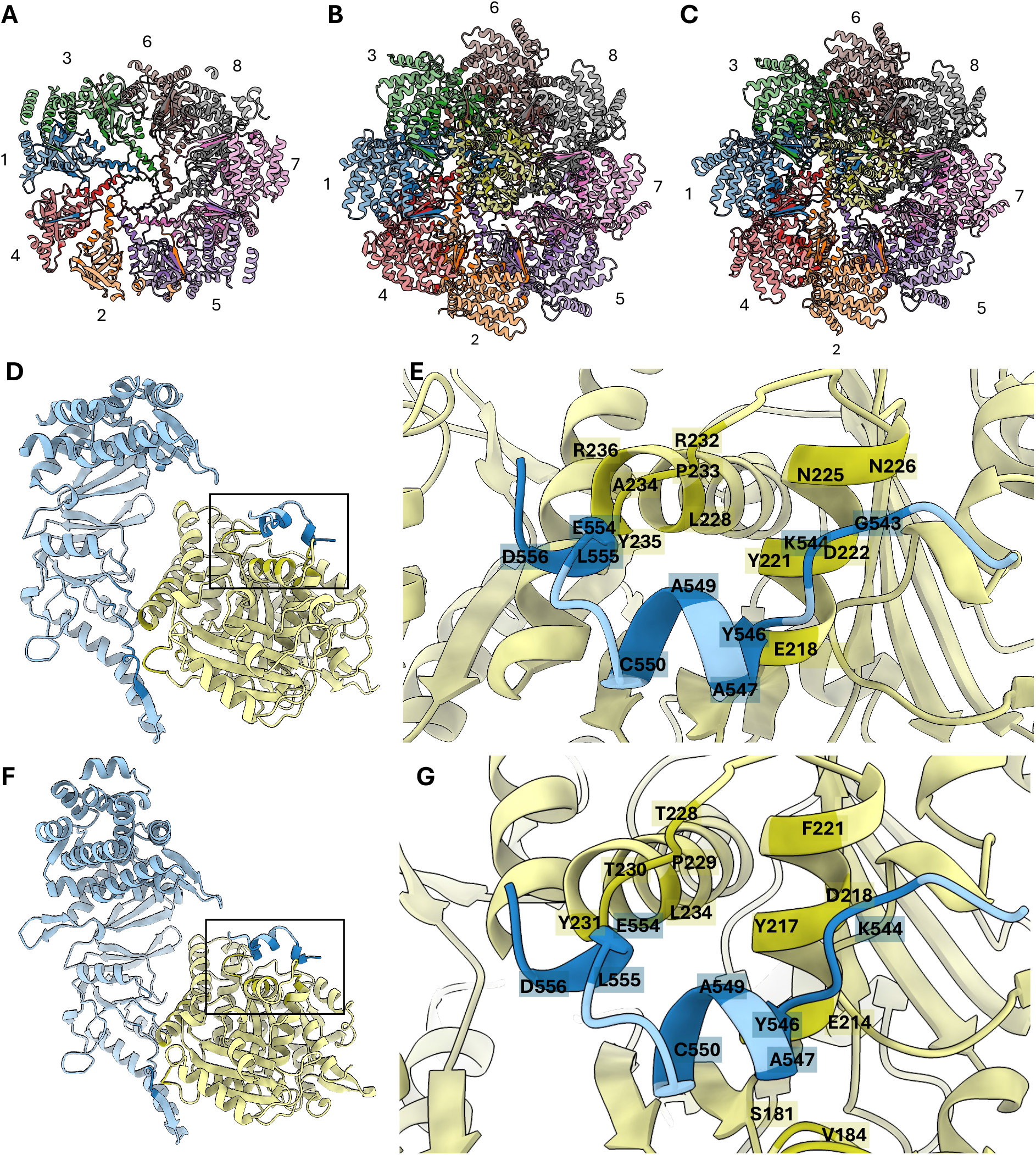
*Drosophila* CCT chaperonin AlphaFold3 predictions in apo form and bound to α- and β-tubulin substrates. **(A–C)** ChimeraX views of the full complexes from LIVIA’s automatically exported ChimeraX scripts; each subunit is colored individually, with LIR shown in the light chain color and cLIR in the dark chain color. CCT subunits are labeled 1–8 (corresponding to CCT1–CCT8). **(A)** Apo CCT. **(B)** CCT bound to α-tubulin (αTub67C). **(C)** CCT bound to β-tubulin (βTub56D). The substrate-bound complexes show markedly more LIR/cLIR-supported structure than the apo complex (mean pTM 0.58 / ipTM 0.55 for apo vs 0.71 / 0.69 for αTub67C and 0.74 / 0.73 for βTub56D, across 5 models). **(D, E)** Detailed view of the CCT1–αTub67C pair from the substrate-bound prediction, rendered with LIVIA’s automatically exported ChimeraX script. CCT1 is shown in light blue and αTub67C in light olive, with non-interacting regions faded; the box in **D** marks the region magnified in **E. (E)** Close-up showing labeled cLIR positions between the CCT1 C-terminal tail and αTub67C. **(F, G)** As in **D, E** but for the CCT1–βTub56D pair. **(F)** Overall dimer view; the box marks the region magnified in **G. (G)** Close-up showing labeled cLIR positions between the CCT1 C-terminal tail and βTub56D.

